# N-Terminal Alkylation of Proteins with Triazole-4-carbaldehyde for Targeted Liposome Engineering

**DOI:** 10.64898/2026.07.31.742023

**Authors:** Lisa Connolly, Akimitsu Okamoto, Neal K. Devaraj, Akira Onoda

## Abstract

A platform method to associate native proteins and peptides onto a liposome membrane using site-specific N-terminal alkylation based on 1*H*-1,2,3-triazole-4-carbaldehyde (TA4C) is developed. The TA4C reagent reacts with the N-terminal α-amino group of native proteins under mild aqueous conditions in a single step, without genetic engineering or protecting group strategies. Equipping TA4C with hexyl and nonyl chains provides a direct handle for tuning the association between the protein and membrane. N-terminal alkylation of green fluorescent protein (GFP) as a model proceeds in high yield (93% for the hexyl group and 70% for the nonyl group), and tethering the N-terminal alkyl group on GFP efficiently associates the protein with the liposomal membrane, as confirmed by confocal laser scanning microscopy and dynamic light scattering. We extended this strategy to an investigation of the GE11 peptide, a ligand for the epidermal growth factor receptor (EGFR). The liposome immobilized with GE11 peptide possessing an N-terminal alkyl group enables active targeting with EGFR-overexpressing A431 cells.

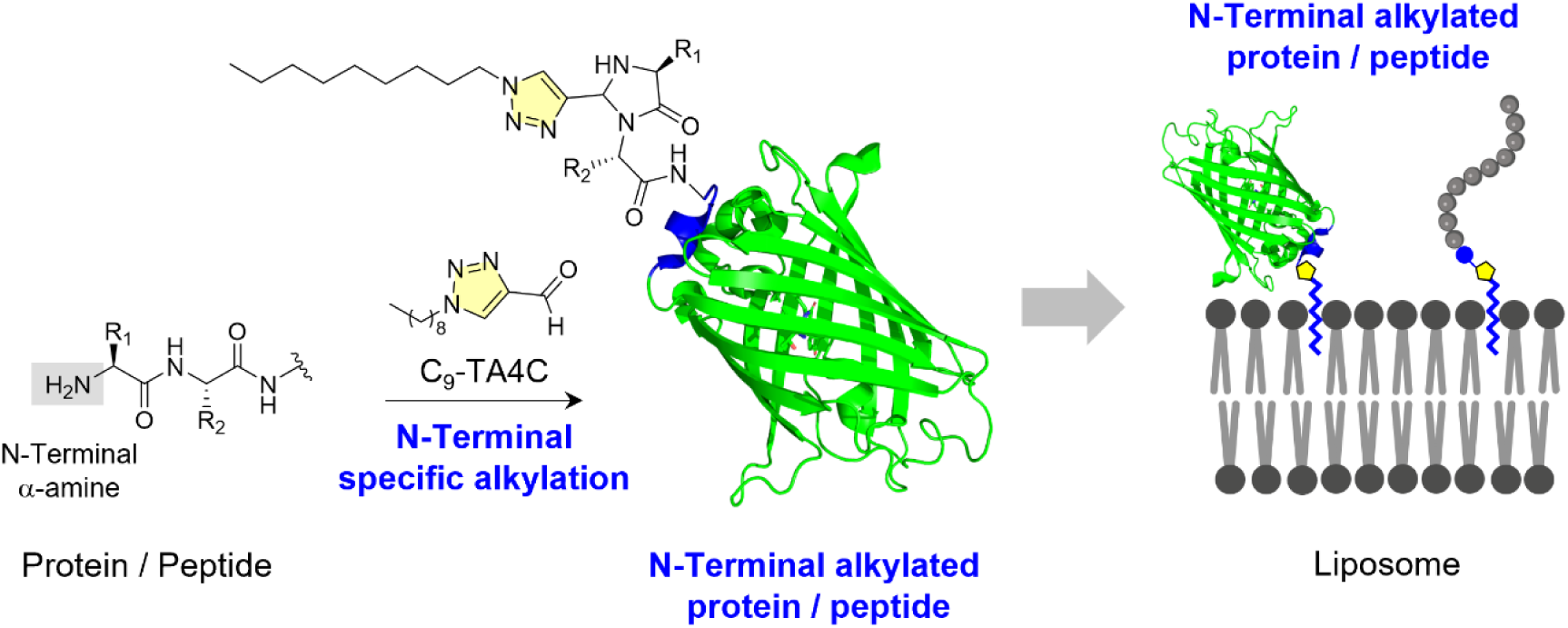

## INTRODUCTION

Enzymatic lipidation governs the membrane association of proteins in living systems. Anchoring systems that use N-myristoylation, S-palmitoylation, prenylation, and glycosylphosphatidylinositol (GPI) each include attachment of a hydrophobic moiety at a defined residue and thereby convert a soluble protein into a membrane-associated species.^1–3^ These modifications control whether a protein resides at a membrane, and where and to which organelle it accumulates within the cell. Biophysical studies of lipidated Ras peptides have demonstrated that the amounts and chemical structures of the attached lipids determine the depth of membrane insertion and the residence time at the lipid bilayer.^4^ The reversibility of the attachment of palmitoylation on the protein was shown to regulate dynamic redistribution.^5^ The covalent attachment of a lipid entity at a single residue plays a key role in maintaining protein functionality associated with membranes.

Considerable effort has thus been devoted to installing hydrophobic anchors site-specifically onto proteins in an artificial manner by enzymatic and chemical methods.^6–9^ Microbial transglutaminase installs lipid anchors on proteins site-specifically,^10^ and the length of the appended acyl chain governs membrane anchoring and the resulting cell selectivity.^11^ Sortase A appends lipid moieties at an engineered LPXTG motif.^12^ Each of these reactions proceeds with high site-selectivity by carrying the requisite recognition sequence. Chemical and semisynthetic routes relax this constraint and allow the anchor itself to be varied independently of the protein sequence.^8,9^ Bioorganic synthesis of lipid-modified Ras was found to enable the reconstitution of signaling events on model membranes.^13^ Native chemical ligation and click chemistry of a synthetic lipid anchor generates a lipidated green fluorescent protein that can be inserted stably into supported bilayers,^14,15^ and the diacylation of peptides has recently yielded functional drug-carrying vesicles.^16^ Most bioconjugation reagents target natural amino acids such as Lys ε-amino or Cys thiol groups to generate lipidated protein in a facile manner.^17,18^ The conjugation with Lys produces a heterogeneous mixture and Cys modification requires the presence of a unique residue within a whole protein.

The N-terminal α-amino group occupies a privileged position among the reactive handles available on native proteins, since it constitutes a single chemically distinct site per polypeptide chain.^19–21^ The p*K*_a_ of an N-terminal α-amine generally lies below that of lysine ε-amino groups,^22^ which allows reaction conditions to be specifically used with the α-amino group under neutral pH conditions. In addition, as the N-terminus is positioned at the flexible end of a protein chain, the N-terminal modification generally avoids interference with the folding and function of the protein.^23^ This advantage has led to the discovery of various types of N-terminus-specific modification reactions using *N*-phenyl-*N*-acetovinylsulfonamide via aza-Michael addition,^24^ and *o*-aminophenols or *o*-catechols that react with N-terminal proline.^25^ Importantly, 2-pyridinecarbaldehyde derivatives are specifically linked to an N-terminal amino group forming an imidazolidinone ring.^20^ A copper-mediated cycloaddition between maleimides and 2-pyridinecarboxaldehyde has recently been shown to be capable of installing diverse functional groups at the N-terminus.^26–28^ Each of these strategies demonstrate the growing repertoire of chemical tools for N-terminal labeling and immobilization.

Our group previously developed 1*H*-1,2,3-triazole-4-carbaldehyde (TA4C), a reagent that reacts with the N-terminal α-amino group of native proteins in a single step under mild aqueous conditions and gives a stable 4-imidazolidinone adduct without perturbing protein structure or function.^29–31^ A hydrophobic tether installed at the N-terminus of a cutinase was further found to enhance the adsorption of the enzyme to a solid poly(ethylene terephthalate).^32,33^ This result demonstrated a broader potential of N-terminal hydrophobic modification to direct proteins toward hydrophobic interfaces. Here we report TA4C reagents tethering alkyl chains and their application to the immobilization of proteins on liposomal membranes. Green fluorescent protein (GFP) was used as a reporter, and we investigated the membrane association of GFP tethered to a hydrophobic N-terminal tail using liposomes as models (Figure 1). Extension of this chemistry to GE11, a peptide ligand for the epidermal growth factor receptor,^34^ yields targeted liposomes modified with GE11 peptide by membrane anchoring with an N-terminal hydrophobic tail. The liposomes decorated with N-terminally alkylated GE11 peptides enable active targeting toward EGFR-overexpressing A431 cells.

**Figure 1.**
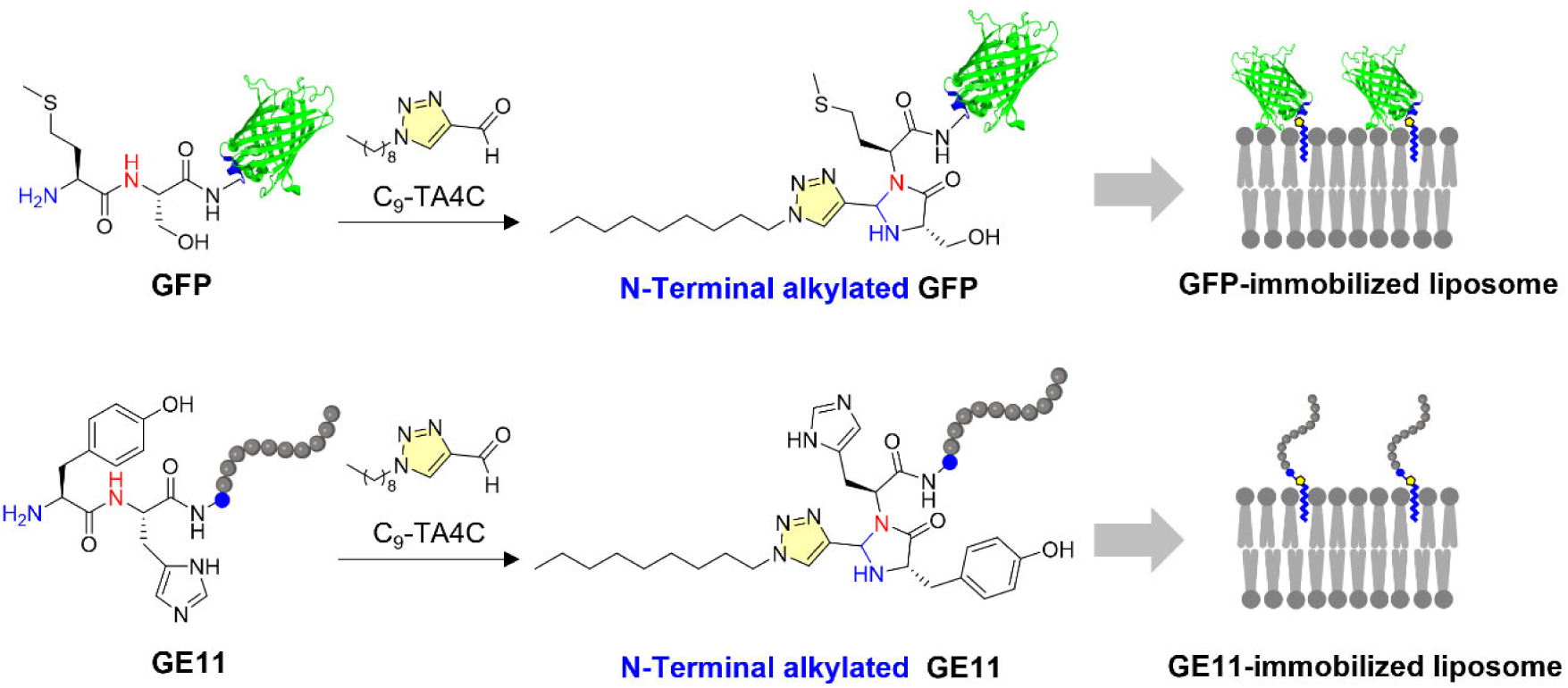
Schematic illustration of N-terminal specific alkylation of protein and peptides by 1*H*-1,2,3-triazole-4-carbaldehyde (TA4C). The N-terminal alkylated protein and peptide are immobilized on liposome.

## RESULTS AND DISCUSSION

### N-Terminal Alkylation of GFP Using C_6_-TA4C, C_9_-TA4C, and C_12_-TA4C

Three versions of the N-terminal specific modification regent TA4C were prepared with hexyl, nonyl, and dodecyl groups according to a previously reported method.^29,32^ In brief, the corresponding alkyl amine compounds were reacted with p-nitrophenyl TA4C to give TA4C derivatives. GFP was reacted with C_6_-TA4C, C_9_-TA4C, and C_12_-TA4C derivatives in 100 mM potassium phosphate buffer (pH 7.5) containing 5% (v/v) of DMSO at 37 °C for 16 h. The reaction proceeds via condensation of the TA4C aldehyde with the N-terminal α-amino group of GFP, forming a stable 4-imidazolidinone ring that incorporates the N-terminal Met residue and the second Ser residue (Figure 2). The modification yields of C_6_-TA4C and C_9_-TA4C were determined by LC-ESI-TOF MS (Figure 2). The yields of C_6_-GFP (obs *m*/*z* 28,206) and C_9_-GFP (obs *m*/*z* 28,246) were 93% and 70%, respectively. C_12_-TA4C did not give detectable conversion under these conditions due to low aqueous solubility of C_12_-TA4C, although we observed conversion in a solution with a higher DMSO concentration of 80% (v/v). The fluorescence measurement of GFPs tethering an N-terminal alkyl chain confirms that the N-terminal modification does not affect the fluorescence properties, with the fluorescence maxima observed at 512 nm (*λ*_ex_ = 488 nm) (Figure S1).

**Figure 2.**
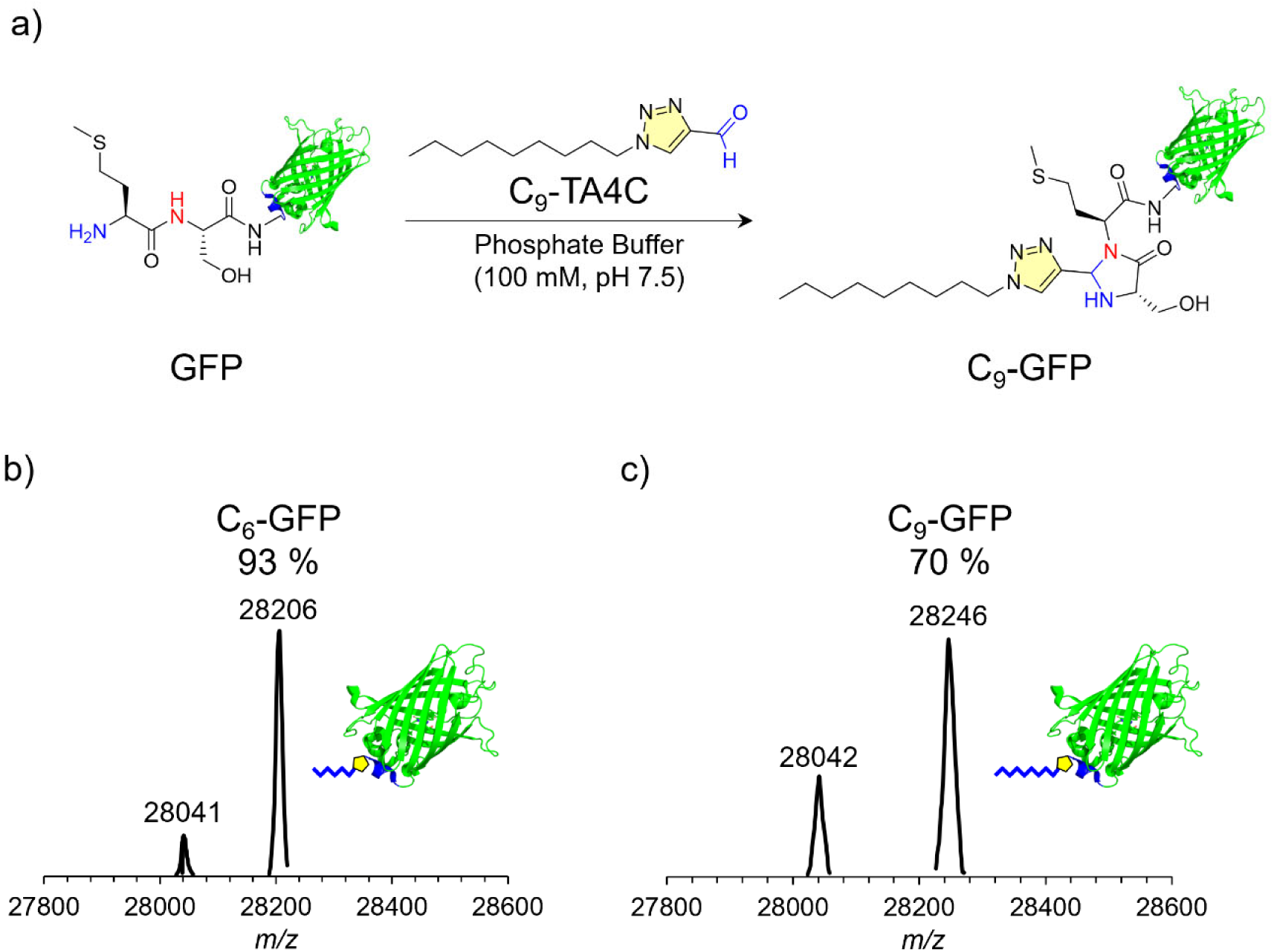
a) N-Terminal specific alkylation of GFP using TA4C. Deconvoluted ESI-TOF MS spectra of b) C_6_-GFP and c) C_9_-GFP.

### Immobilization of GFP Tethering N-Terminal Alkyl Chain onto Liposomes

Liposomes with average diameter of ca. 5 μm were prepared for visualization using fluorescence microscopy. Liposomes containing a lipid linked with rhodamine were prepared by hydration and visualized by confocal fluorescence microscopy (Figure S2). The clearly observed giant unilamellar vesicles were found to have an average diameter of 4.65±1.8 μm. The liposomes were then incubated with C_6_-GFP, C_9_-GFP, and GFP, and visualized (Figure 3). In the case of the liposome treated with C_6_-GFP, the GFP signal is clearly localized on the liposome membrane (Figure 3a). The average size of the liposomes is slightly smaller before the addition of C_6_-GFP. In the case of the liposomes treated with C_9_-GFP, the GFP signal is also localized on the liposome membrane (Figure 3b). We observed aggregated liposomes, suggesting the possibility of perturbation of the C_9_ chain linked with GFP toward the feature of the membrane. In stark contrast, GFP without modification is diffused in the solution, and the interaction between GFP and the membrane is not observed (Figure 3c). These results clearly demonstrate that GFPs tethering an N-terminal alkyl chain are immobilized onto the lipid bilayer of the liposomes driven by the hydrophobic tail.

**Figure 3.**
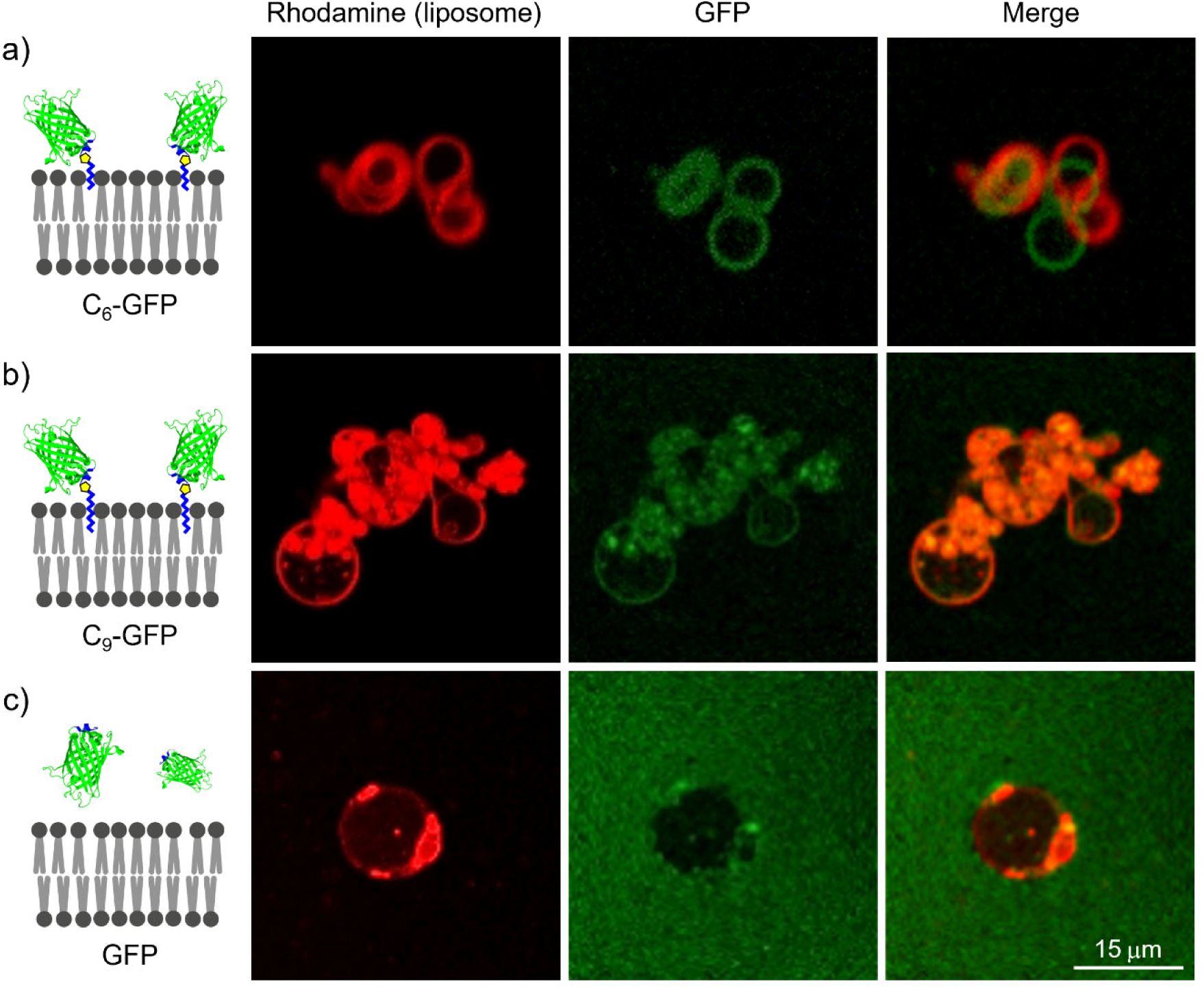
Confocal Laser Scanning Microscopy (CLSM) images of liposomes (size, 4.65±1.8 μm) treated with (a) C_6_-GFP, (b) C_9_-GFP, and (c) unmodified GFP.

In a cell targeting experiment, liposomes with a smaller size were prepared and tested for the immobilization of GFP with an N-terminal alkyl group. Because a size less than 150 nm is required for internalization into the cell, the liposomes were produced by extrusion through a 100 nm nominal pore-size polycarbonate membrane, and the average size was analyzed by dynamic light scattering (DLS) measurement. The average hydrodynamic diameter of the liposomes was found to be 132±10 nm with a narrow and unimodal size distribution, and the measured ζ potential was −33±1 mV. The size and ζ potential values of the liposomes were analyzed after treatment with C_6_-GFP, C_9_-GFP, and GFP. The liposomes treated with C_6_-GFP exhibited a smaller size of 120±8 nm. This trend is in accordance with the result that we observed in fluorescence microscopy using larger liposomes with (Figure 3a). The liposomes treated with C_9_-GFP exhibited an increased size of 154±11 nm. The result is also consistent with the observation of aggregation of the larger liposomes (Figure 3b). A change in the size (132±10 nm) after the addition of GFP was not observed. In addition, the ζ-potential values are also slightly changed only in the case of addition of C_6_-GFP (−29±2 mV) and C_9_-GFP (−28±3 mV). The addition of GFP does not change the ζ-potential (−31 ± 2 mV). To directly evaluate protein immobilization on the surfaces of liposomes, rhodamine-labeled liposomes incubated with C_6_-GFP, C_9_-GFP, or GFP were analyzed by confocal fluorescence microscopy (Figure 4). As expected, liposomes treated with C_6_-GFP and C_9_-GFP show GFP signals which are significantly co-localized with the red rhodamine signal of liposomes (Figures 4a and 4b). In contrast, liposomes treated with GFP do not show the co-localized signals (Figure 4c). The DLS and imaging results clearly support the finding that both C_6_-GFP and C_9_-GFP are capable of being immobilized on the membrane of liposomes with average diameters less than 150 μm that can be internalized into the cells.

**Figure 4.**
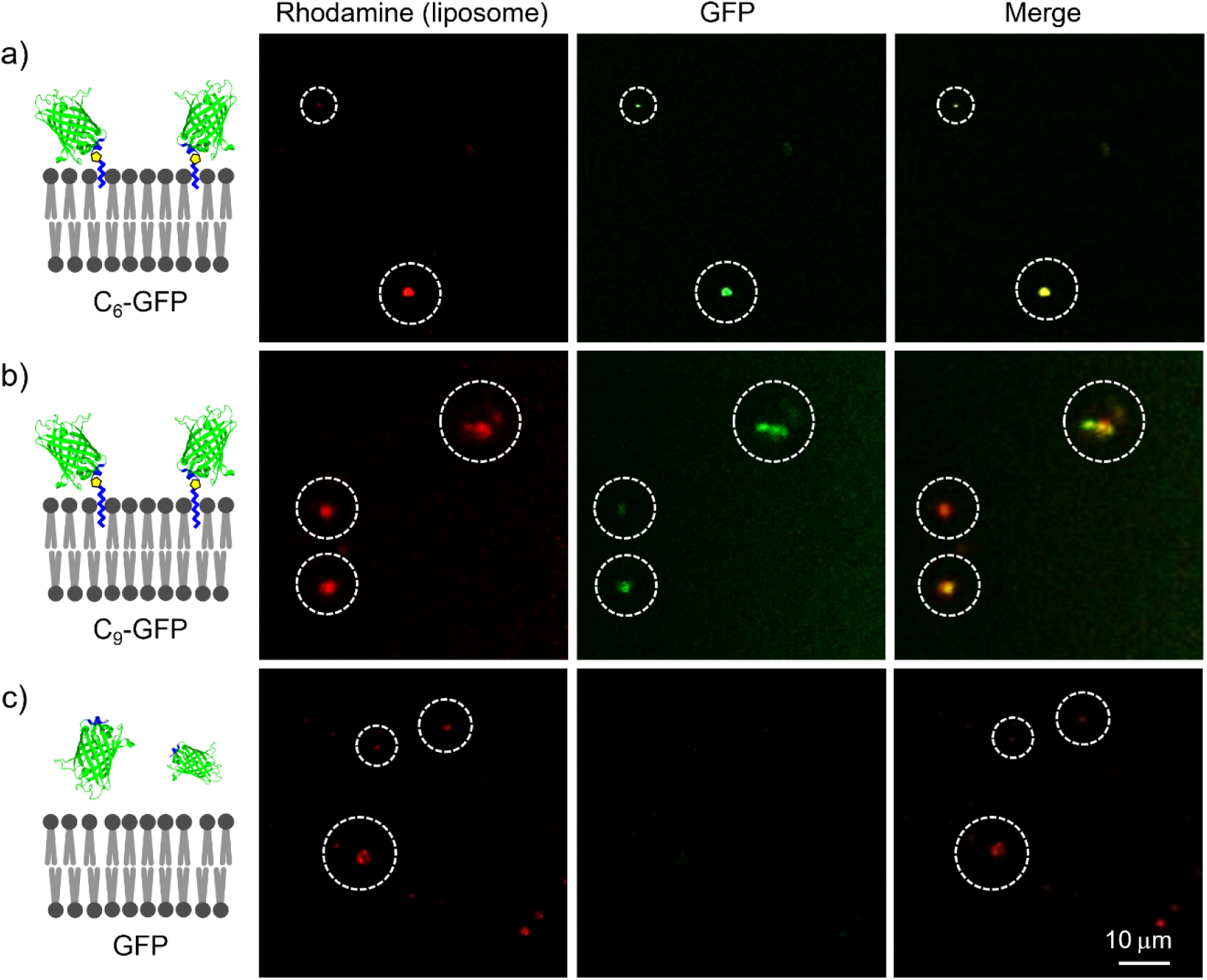
CLSM images of liposomes (size, 132±10 nm) treated with (a) C_6_-GFP, (b) C_9_-GFP and (c) GFP.

To quantitatively assess the immobilization efficiency, the supernatant fluorescence was measured after centrifugation of C_n_-GFP immobilized liposome (6,000 rpm, 4 min, 4 °C). Approximately 50% of the fluorescence was retained in the supernatant for both C₆-GFP and C₉-GFP, while nearly 100% was retained for unmodified GFP, suggesting that approximately half of the alkyl-modified GFP molecules were immobilized on the liposomal membrane.

### Active Cell Targeting of Liposomes Functionalized with GE11 Containing an N-Terminal Alkyl Group

Encouraged by the successful results on surface immobilization of liposomes by proteins tethering an alkyl chain at the N-terminus, we next investigated whether the liposomes immobilized with the ligands are capable of active targeting on the cell surface. GE11 specifically binds to epidermal growth factor receptor (EGFR).^34–38^ First, the N-terminus of the GE11 peptide was modified using C_6_-TA4C and C_9_-TA4C. The GE11 peptides tethering an alkyl chain were incorporated into the liposomes labeled with rhodamine. A431 cells overexpressing EGFR were used to evaluate the potential for active targeting of liposomes. The A431 cells were seeded with 1.0 × 10⁵ cells per well in 96-well plates and cultured for 24 h before the treatment. Liposomes were supplemented to provide a lipid concentration of 160 μM in 100 μL per well, a condition that produced consistently strong and reproducible fluorescence signals in preliminary experiments. Four formulations were prepared, including (1) liposomes modified with C_6_-GE11, (2) liposomes modified with C_9_-GE11, (3) liposomes mixed GE11, and (4) liposomes without any treatment as negative controls. After addition of the liposomes to the cell with incubation for 1 h at 37 °C, the cells were imaged directly by confocal laser scanning microscopy without washing to capture total cellular association. We found clear spots of rhodamine-labeled liposomes within the cells treated with liposomes containing C_6_-GE11 and C_9_-GE11 (Figures 5a and 5b). In contrast, the number of spots within the cell is clearly reduced in liposomes mixed with GE11 and plain liposomes without treatment (Figures 5c, 5d and sS4). These results indicate that N-terminal modification of GE11 with an alkyl chain harnesses the potential of active targeting of liposomes mediated by the specific interaction between GE11 ligands and EGFR on the surface of A431 cells. These results clearly demonstrate that the GE11 peptide modified on the liposome membrane contributes to the robust engagement of EGFR on the surface of A431 cells.

**Figure 5.**
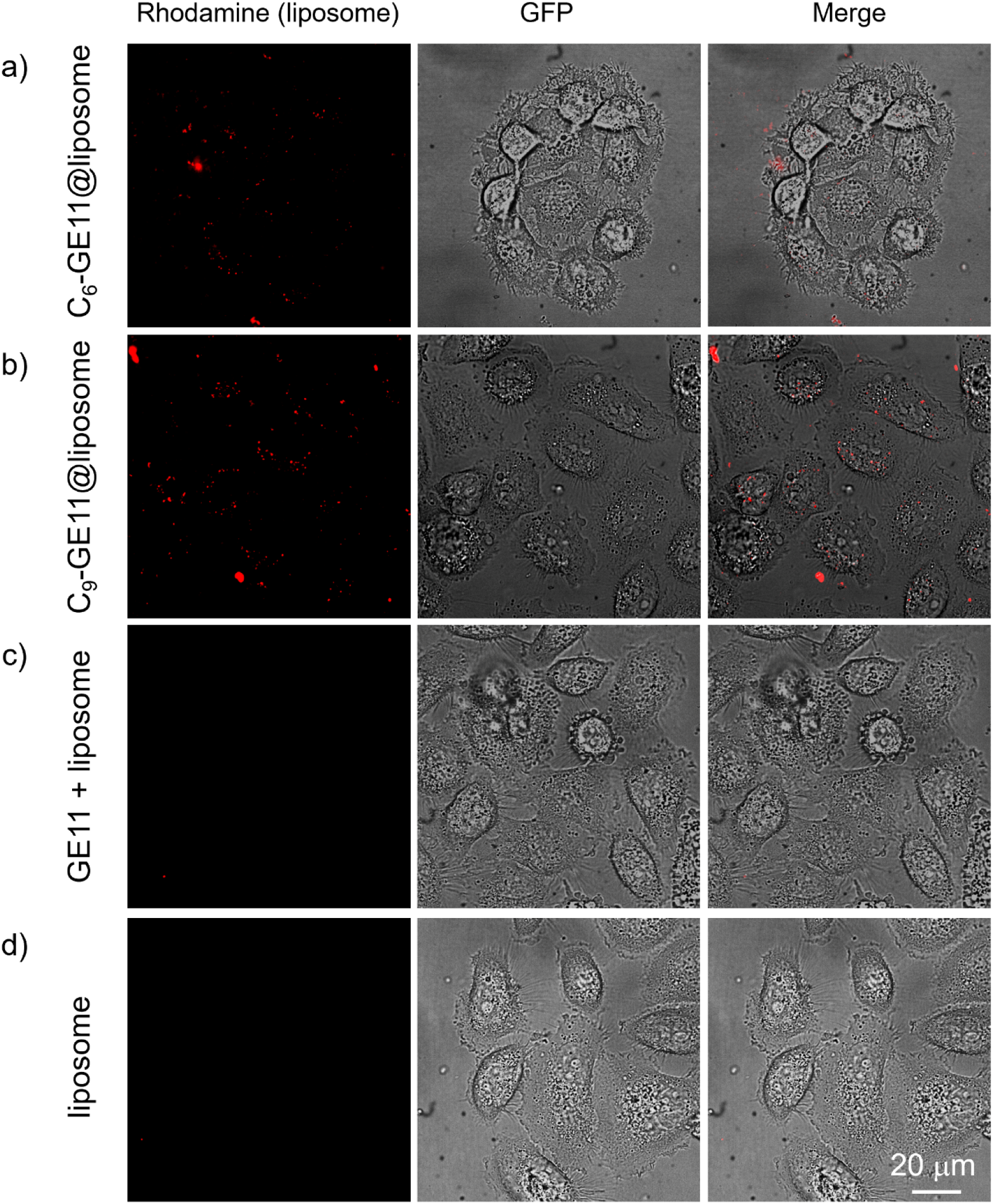
CLSM images of A431 cells after the addition of (a) liposomes modified with C_6_-GE11, (b) liposomes modified with C_9_-GE11, (c) liposomes mixed with GE11, and (d) unmodified liposomes.

## CONCLUSIONS

This study demonstrates that N-terminal alkylation using 1*H*-1,2,3-triazole-4-carbaldehyde-(TA4C) provides a site-specific bioconjugation process for anchoring native proteins to liposomal membranes without genetic engineering. N-Terminal alkylation of GFP with hexyl- and nonyl-TA4C proceeds in 93% and 70% yield, respectively, under mild aqueous conditions, and the resulting conjugates efficiently associate with liposomal membranes, as confirmed by confocal fluorescence microscopy and dynamic light scattering. Extension of this chemistry to the EGFR-targeting peptide GE11 yields liposomes that associated selectively with EGFR-overexpressing A431 cells. These results establish TA4C-based N-terminal alkylation as a simple and versatile platform for constructing protein-liposome interfaces and targeted liposome-based delivery systems.

## EXPERIMENTAL SECTION

### Materials

Reagents and solvents were used as received without purification unless otherwise noted. All chemicals and organic solvents utilized for synthesis in this study were purchased from FUJIFILM Wako Pure Chemical Corporation, Tokyo Chemical Industry Co., Ltd. (TCI), Sigma-Aldrich Japan, and Aldrich Chemical Co, Inc. The triazole compounds used as precursors were synthesized with reference to previous reports.^29,32^ Palmitoyl oleoyl phosphatidylcholine (POPC), cholesterol (Chol), and 1,2-dipalmitoyl-*sn*-glycero-3-[phospho-*rac*-(1-glycerol)] (sodium salt) (DPPG), 1,2-dioleoyl-*sn*-glycero-3-phosphoethanolamine (DOPE), Rho-DHPE (N-(lissamine rhodamine B sulfonyl)-1,2-dipalmitoyl-*sn*-glycero-3-phosphoethanolamine, triethylammonium salt) were obtained from FUJIFILM Wako. Ultrapure water was demineralized using a Merck Milli-Q IQ 7005 system.

### Synthesis of C_6_-TA4C, C_9_-TA4C, and C_12_-TA4C

C_6_-TA4C, C_9_-TA4C, and C_12_-TA4C were synthesized from hexylamine, nonylamine, dodecylamine, and *p*-nitrophenyl TA4C according to a previously reported method.^29,32^

### N-terminus modification of GFP using C_n_-TA4C

GFP in potassium phosphate buffer (100 μM, 25 μL, final concentration: 50 μM) was diluted with potassium phosphate buffer (100 mM, pH 7.5, 22.5 μL). To the resulting solution, TA4C derivatives in DMSO (200 mM, 2.5 μL, final concentration: 10 mM) were added, and the mixture was incubated at 37 °C for 16 h. The protein was then purified using a Low Protein Binding Durapore Membrane (0.22 μm). Modified GFP preparations were analyzed by LC-MS (Figure 2).

### Characterization of C_6_-GFP and C_9_-GFP

Electrospray ionization time-of-flight mass spectrometry (ESI-TOF MS) spectra were acquired using a Bruker MicrOTOF II HE. Modification of GFP was monitored by LC-MS using a gradient of CH_3_CN containing 0.1% formic acid. LC was conducted with a Poroshell 300SB-C3 column (Agilent Technologies) at 25 °C with a flow rate of 0.2 mL min^−1^. The conversion yield was evaluated from the peak intensity in ESI-TOF MS.

### Preparation of Liposomes

POPC (0.30 mg), cholesterol (0.15 mg), Rhodamine-DHPE (1.3 μg), and 460 μL of chloroform: methanol mixture solution (2: 1) were added into a 5 mL pear-shaped flask. The mixture was stirred for 1 hour on a stirrer to form a homogeneous solution. After organic solvent was removed *in vacuo*, a homogeneous phospholipid film was hydrated with 2 mL of phosphate buffer (100 mM, pH 7.5), and the sample was stored for 24 h at 4 °C. The resulting mixture was briefly ultrasonicated (1–2 s) to form large multilamellar vesicles (MLVs). (PC: Cholesterol:Rho-DHPE = 1000 : 1000 : 2.5) Liposomes with a smaller size were produced by passing the MLVs through the extruder for 11 times.

### Characterization of Liposomes

The particle size, ζ potential, and PDI were measured by dynamic light scattering with Malvern Zeta particle size analyzer (Nano-ZS, Worcestershire, UK). Liposomes with average diameter of 4.4 μm were prepared by hydration without extrusion and visualized by confocal fluorescence microscopy. Spherical and giant unilamellar vesicles suitable for fluorescence imaging were prepared. Small liposomes were produced by extrusion through a 100 nm nominal pore-size polycarbonate membrane and analyzed by dynamic light scattering (DLS). The measurements indicated an average hydrodynamic diameter of 132±10 nm, with a narrow, unimodal size distribution.

### Immobilization of C_6_-GFP and C_9_-GFP onto Liposomes

Liposomes with a lipid concentration: of 0.2 mM (100 μL) were mixed with the solution of GFP (1 μM, 10 μL) and the mixed solution was incubated at 25 °C for 4 h. The samples were stored at 4 °C and used for confocal microscopy experiments.

### Cells Culture and Assays

A431 cells (human epidermoid carcinoma cells) were cultured in Gibco Dulbecco’s Modified Eagle Medium (DMEM) containing L-glutamine and sodium pyruvate, supplemented with 10% (v/v) fetal bovine serum (FBS), 1× Gibco GlutaMAX, and 1× Gibco penicillin–streptomycin, at 37 °C in a humidified atmosphere containing 5% CO₂..

### Cellular Binding and Uptake Studies of GE11-modified liposomes

EGFR-overexpressing A431 cells (human epidermoid carcinoma) were seeded in 96-well plates at a density of 1.0 × 10⁵ cells per well and cultured for 24 h before performing the experiments. Cells were then incubated with rhodamine-labeled liposomes at a lipid concentration of 160 μM in 100 μL per well. Four liposome formulations, (1) liposomes modified with C_6_-GE11, (2) liposomes modified with C_9_-GE11, (3) liposomes mixed with GE11, and (4) unmodified liposomes as negative controls, were prepared. The cells were incubated with liposomes for 1 h at 37 °C in Hanks’ balanced salt solution (HBSS). The cells were imaged directly by confocal laser scanning microscopy (Zeiss Z1 Spinning Disk) without washing to evaluate the association of liposomes with the cell surface.

## ASSOCIATED CONTENT

### Supporting Information

The Supporting Information is available free of charge on the ACS Publications website. Synthesis TA4C synthesis, protein modification, preparation of liposome, and fluorescence imaging data (PDF).

## ACKNOWLEDGMENTS

This work was supported by JSPS KAKENHI Grant Number JP24H02213 in Transformative Research Areas (A) JP24A202 Integrated Science of Synthesis by Chemical Structure Reprogramming (SReP) and JP24K01533, SATREPS Project “Recovering High-Value Bioproducts for Sustainable Fisheries in Chile (ReBiS)” funded by JST/JICA (Grant Number JPMJSA2206), and JST Grant Number JPMJTM20A8 and JPMJTM20JE to A.O. L.C. acknowledges support from Hokkaido University EXEX Doctoral Fellowship (JST SPRING, Grant Number JPMJSP2119). We acknowledge Prof. Katsuaki Konishi at Faculty of Environmental Earth Science, Hokkaido University, for DLS and ζ potential measurements and fluorescence spectroscopy, the Nikon Imaging Center and Advanced Bioimaging Support Platform (Abis) at Hokkaido University supported by JSPS KAKENHI Grant Number JP22H04926 for confocal fluorescence microscopy experiments, and Evan McCormack at the University of California San Diego for his invaluable support throughout the cell-based experiments, including cell culture, confocal fluorescence microscopy, and many other aspects of the experimental work.

---

**Figure S1.**
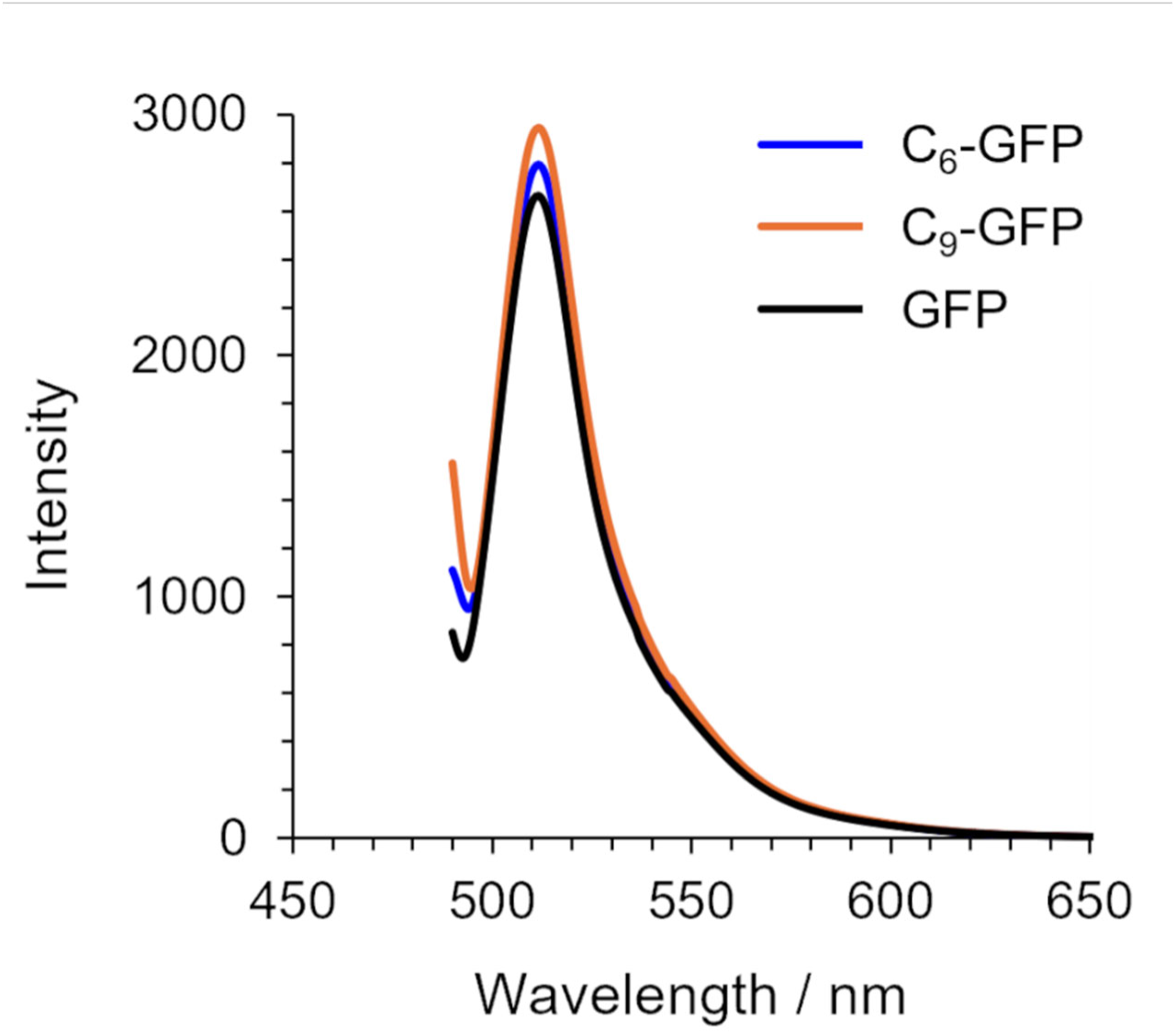
Fluorescence spectra of C_6_-GFP (red), C_9_-GFP (blue), and modified GFP (black).

**Figure S2.**
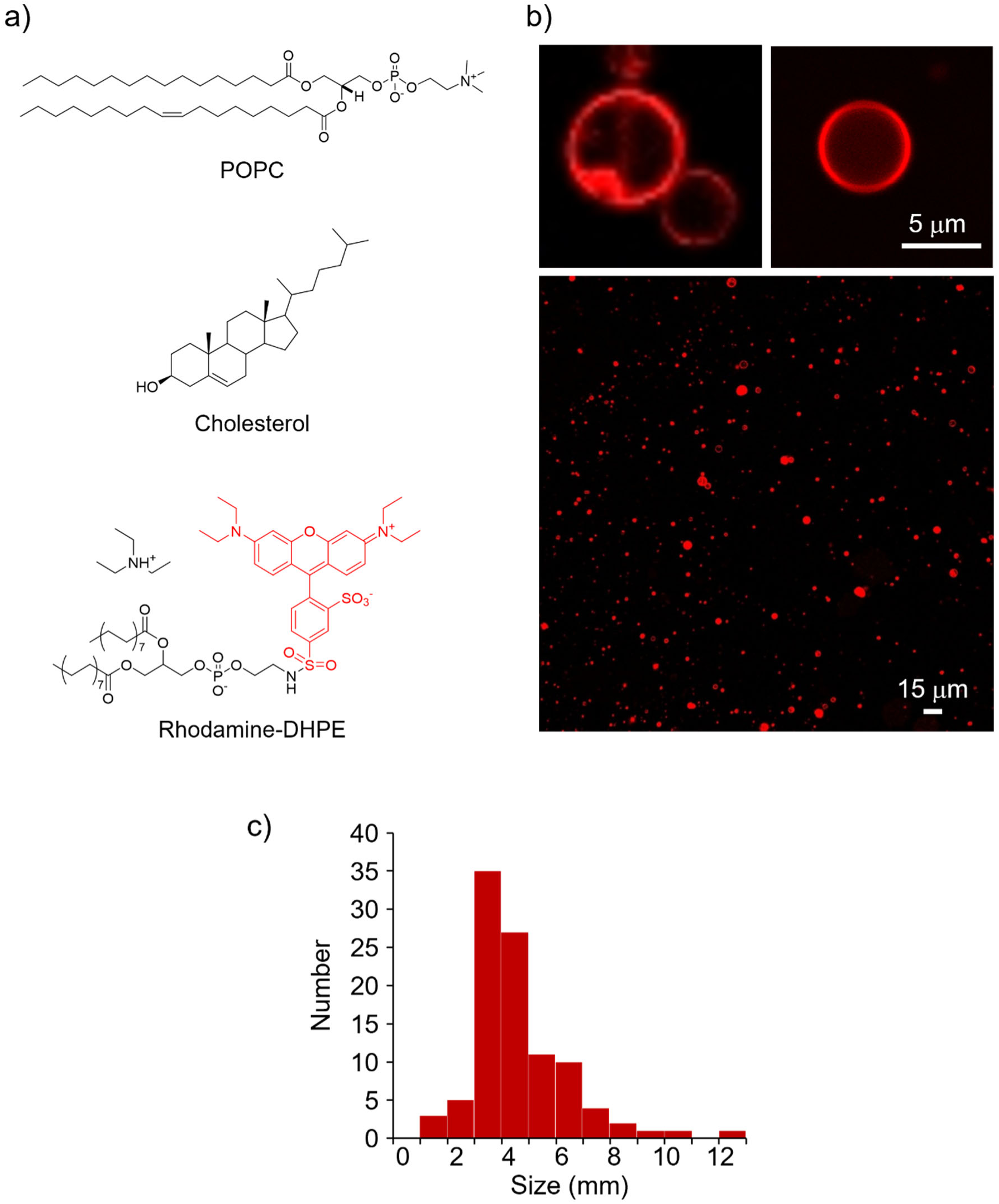
(a) Composition of liposome. (b) CLSM images of liposomes of zoomed and large view. (c) Size distribution of the liposomes.

**Figure S3.**
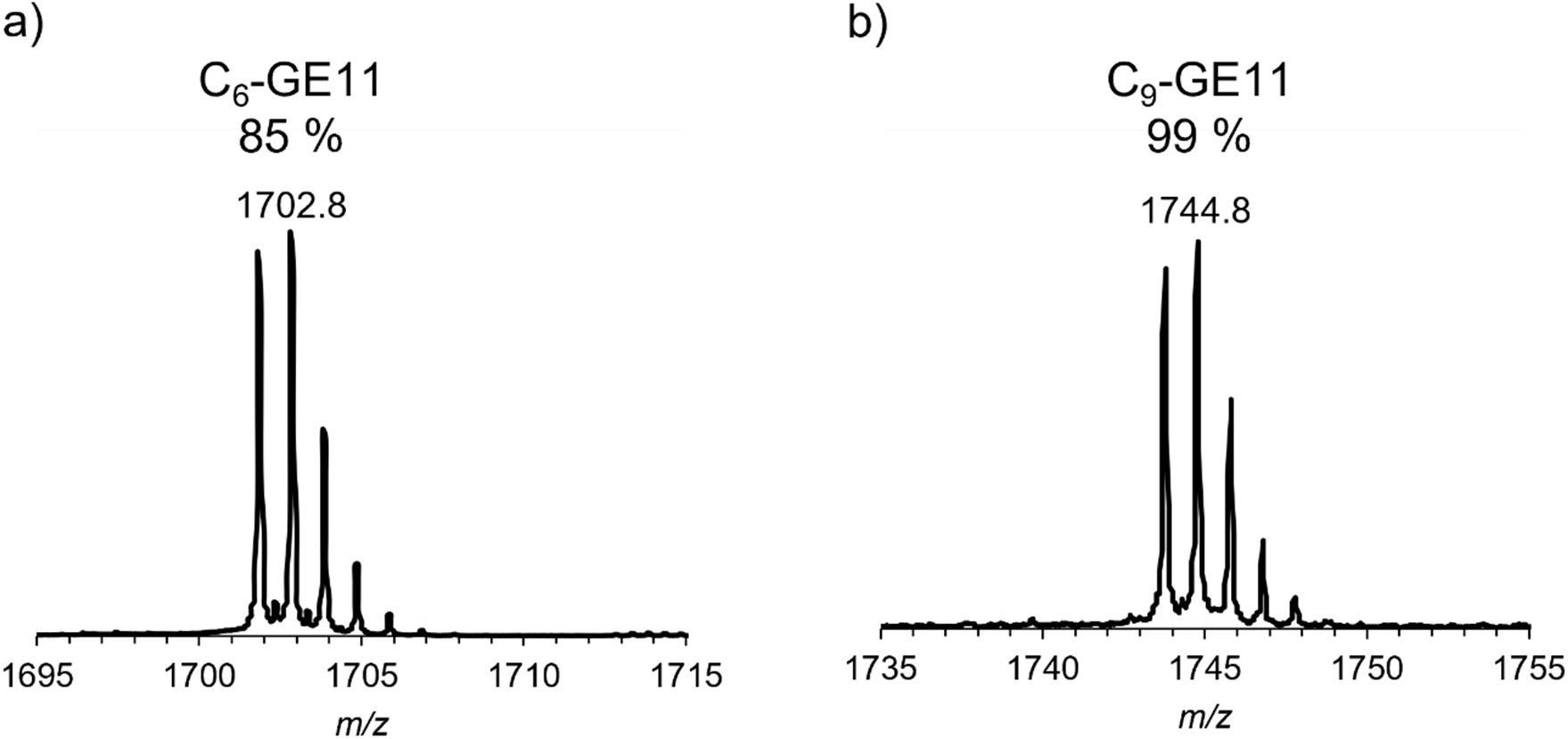
Deconvoluted ESI-TOF MS spectra of (a) C_6_-GE11 and (b) C_9_-GE11

**Figure S4.**
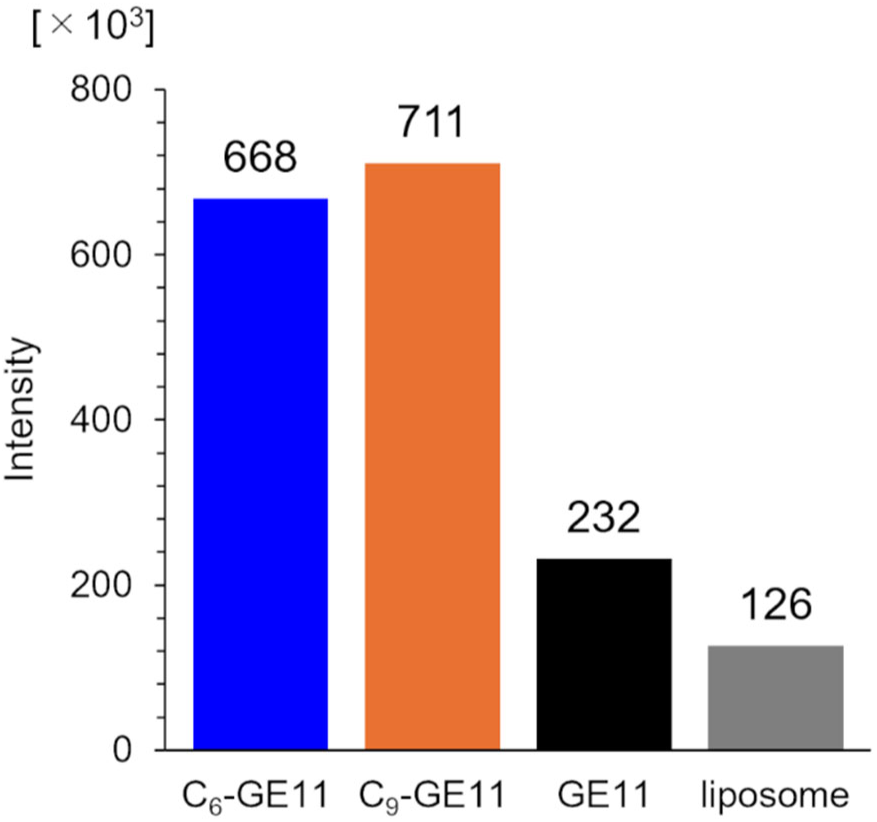
Fluorescent intensity of CLSM images of A431 cells after the addition of liposomes modified with C_6_-GE11, liposomes modified with C_9_-GE11, liposomes mixed with GE11, and unmodified liposomes.

## Notes

### Competing Interest Statement

The authors have declared no competing interest.

## REFERENCES

(1) Ding, W.; Gu, J.; Xu, W.; Wu, J.; Huang, Y.; Zhang, S.; Lin, S. The Biosynthesis and Applications of Protein Lipidation. Chem. Rev. 2024, 124 (21), 12176–12212. DOI: 10.1021/acs.chemrev.4c00419.

(2) Jiang, H.; Zhang, X.; Chen, X.; Aramsangtienchai, P.; Tong, Z.; Lin, H. Protein Lipidation: Occurrence, Mechanisms, Biological Functions, and Enabling Technologies. Chem. Rev. 2018, 118 (3), 919–988. DOI: 10.1021/acs.chemrev.6b00750.

(3) Gelb, M. H.; Brunsveld, L.; Hrycyna, C. A.; Michaelis, S.; Tamanoi, F.; Van Voorhis, W. C.; Waldmann, H. Therapeutic intervention based on protein prenylation and associated modifications. Nat. Chem. Biol. 2006, 2 (10), 518–528. DOI: 10.1038/nchembio818.

(4) Huster, D.; Vogel, A.; Katzka, C.; Scheidt, H. A.; Binder, H.; Dante, S.; Gutberlet, T.; Zschörnig, O.; Waldmann, H.; Arnold, K. Membrane Insertion of a Lipidated Ras Peptide Studied by FTIR, Solid-State NMR, and Neutron Diffraction Spectroscopy. J. Am. Chem. Soc. 2003, 125 (14), 4070–4079. DOI: 10.1021/ja0289245.

(5) Rocks, O.; Peyker, A.; Kahms, M.; Verveer, P. J.; Koerner, C.; Lumbierres, M.; Kuhlmann, J.; Waldmann, H.; Wittinghofer, A.; Bastiaens, P. I. H. An Acylation Cycle Regulates Localization and Activity of Palmitoylated Ras Isoforms. Science 2005, 307 (5716), 1746–1752. DOI: 10.1126/science.1105654.

(6) Takahara, M.; Kamiya, N. Synthetic Strategies for Artificial Lipidation of Functional Proteins. Chem. Eur. J. 2020, 26 (21), 4645–4655. DOI: 10.1002/chem.201904568.

(7) Zhang, Y.; Park, K.-Y.; Suazo, K. F.; Distefano, M. D. Recent progress in enzymatic protein labelling techniques and their applications. Chem. Soc. Rev. 2018, 47 (24), 9106–9136. DOI: 10.1039/c8cs00537k.

(8) Hanna, C. C.; Kriegesmann, J.; Dowman, L. J.; Becker, C. F. W.; Payne, R. J. Chemical Synthesis and Semisynthesis of Lipidated Proteins. Angew. Chem. Int. Ed. 2022, 61 (15), e202111266. DOI: 10.1002/anie.202111266.

(9) Mejuch, T.; Waldmann, H. Synthesis of Lipidated Proteins. Bioconjugate Chem. 2016, 27 (8), 1771–1783. DOI: 10.1021/acs.bioconjchem.6b00261.

(10) Abe, H.; Goto, M.; Kamiya, N. Protein Lipidation Catalyzed by Microbial Transglutaminase. Chem. Eur. J. 2011, 17 (50), 14004–14008. DOI: 10.1002/chem.201102121.

(11) Ogushi, N.; Uchida, K.; Kawaguchi, Y.; Wakabayashi, R.; Goto, M.; Kamiya, N. Lipid chain length-dependent anchoring of artificial lipidated proteins enables cell-selective uptake of extracellular vesicles. Chem. Lett. 2026, 55 (6), upag092. DOI: 10.1093/chemle/upag092.

(12) Antos, J. M.; Miller, G. M.; Grotenbreg, G. M.; Ploegh, H. L. Lipid Modification of Proteins through Sortase-Catalyzed Transpeptidation. J. Am. Chem. Soc. 2008, 130 (48), 16338–16343. DOI: 10.1021/ja806779e.

(13) Bader, B.; Kuhn, K.; Owen, D. J.; Waldmann, H.; Wittinghofer, A.; Kuhlmann, J. Bioorganic synthesis of lipid-modified proteins for the study of signal transduction. Nature 2000, 403 (6766), 223–226. DOI: 10.1038/35003249.

(14) Grogan, M. J.; Kaizuka, Y.; Conrad, R. M.; Groves, J. T.; Bertozzi, C. R. Synthesis of Lipidated Green Fluorescent Protein and Its Incorporation in Supported Lipid Bilayers. J. Am. Chem. Soc. 2005, 127 (41), 14383–14387. DOI: 10.1021/ja052407f.

(15) Charron, G.; Zhang, M. M.; Yount, J. S.; Wilson, J.; Raghavan, A. S.; Shamir, E.; Hang, H. C. Robust Fluorescent Detection of Protein Fatty-Acylation with Chemical Reporters. J. Am. Chem. Soc. 2009, 131 (13), 4967–4975. DOI: 10.1021/ja810122f.

(16) Cho, C. J.; Kang, S.; Pedebos, C.; Khalid, S.; Brea, R. J.; Devaraj, N. K. Diacylation of Peptides Enables the Construction of Functional Vesicles for Drug-Carrying Liposomes. Angew. Chem. Int. Ed. 2025, 64 (20), e202421932. DOI: 10.1002/anie.202421932.

(17) Koniev, O.; Wagner, A. Developments and recent advancements in the field of endogenous amino acid selective bond forming reactions for bioconjugation. Chem. Soc. Rev. 2015, 44 (15), 5495–5551. DOI: 10.1039/c5cs00048c.

(18) Boutureira, O.; Bernardes, G. J. L. Advances in Chemical Protein Modification. Chem. Rev. 2015, 115 (5), 2174–2195. DOI: 10.1021/cr500399p.

(19) Rosen, C. B.; Francis, M. B. Targeting the N terminus for site-selective protein modification. Nat. Chem. Biol. 2017, 13 (7), 697–705. DOI: 10.1038/nchembio.2416.

(20) MacDonald, J. I.; Munch, H. K.; Moore, T.; Francis, M. B. One-step site-specific modification of native proteins with 2-pyridinecarboxyaldehydes. Nat. Chem. Biol. 2015, 11 (5), 326–331. DOI: 10.1038/nchembio.1792.

(21) Gilmore, J. M.; Scheck, R. A.; Esser-Kahn, A. P.; Joshi, N. S.; Francis, M. B. N-Terminal Protein Modification through a Biomimetic Transamination Reaction. Angew. Chem. Int. Ed. 2006, 45 (32), 5307–5311. DOI: 10.1002/anie.200600368.

(22) Sereda, T. J.; Mant, C. T.; Quinn, A. M.; Hodges, R. S. Effect of the α-amino group on peptide retention behaviour in reversed-phase chromatography Determination of the pKa values of the α-amino group of 19 different N-terminal amino acid residues. J. Chromatogr. A 1993, 646 (1), 17–30. DOI: 10.1016/S0021-9673(99)87003-4.

(23) Jacob, E.; Unger, R. A tale of two tails: why are terminal residues of proteins exposed? Bioinformatics 2007, 23 (2), e225–e230. DOI: 10.1093/bioinformatics/btl318.

(24) Huang, R.; Li, Z. H.; Ren, P. L.; Chen, W. Z.; Kuang, Y. Y.; Chen, J. K.; Zhan, Y. X.; Chen, H. L.; Jiang, B. *N*-Phenyl-*N*-aceto-vinylsulfonamides as Efficient and Chemoselective Handles for N-Terminal Modification of Peptides and Proteins. Eur. J. Org. Chem. 2018, 2018 (6), 829–836. DOI: 10.1002/ejoc.201701715.

(25) Obermeyer, A. C.; Jarman, J. B.; Francis, M. B. N-Terminal Modification of Proteins with o-Aminophenols. J. Am. Chem. Soc. 2014, 136 (27), 9572–9579. DOI: 10.1021/ja500728c.

(26) Machida, H.; Kanemoto, K. N-Terminal-Specific Dual Modification of Peptides through Copper-Catalyzed [3+2] Cycloaddition. Angew. Chem. Int. Ed. 2024, 63 (12), e202320012. DOI: 10.1002/anie.202320012.

(27) Hanaya, K.; Taguchi, K.; Wada, Y.; Kawano, M. One-Step Maleimide-Based Dual Functionalization of Protein N-Termini. Angew. Chem. Int. Ed. 2025, 64 (5), e202417134. DOI: 10.1002/anie.202417134.

(28) Hanaya, K.; Yamoto, K.; Taguchi, K.; Matsumoto, K.; Higashibayashi, S.; Sugai, T. Single-Step N-Terminal Modification of Proteins via a Bio-Inspired Copper(II)-Mediated Aldol Reaction. Chem. Eur. J. 2022, 28 (47), e202201677. DOI: 10.1002/chem.202201677.

(29) Onoda, A.; Inoue, N.; Sumiyoshi, E.; Hayashi, T. Triazolecarbaldehyde Reagents for One-Step N-Terminal Protein Modification. ChemBioChem 2020, 21 (9), 1274–1278. DOI: 10.1002/cbic.201900692.

(30) Wang, S.; Sumiyoshi, E.; Inoue, N.; Zheng, X.; Noro, S.-i.; Hayashi, T.; Onoda, A. Ambient Temperature Synthesis of Triazole-4-carbaldehyde Reagent by Dimroth Rearrangement Enabling Facile N-Terminal-specific Modification and Immobilization of Proteins. Bioconjugate Chem. 2026, 37 (5), 899–905. DOI: 10.1021/acs.bioconjchem.5c00583.

(31) Zhang, Y.; Odani, K.; Onoda, A. N-terminal dual functionalization of proteins via copper-mediated [3 + 2] cycloaddition of triazolecarbaldehyde with maleimide. Chem. Lett. 2025, 54 (12), upaf221. DOI: 10.1093/chemle/upaf221.

(32) Shuvo, M. S. R.; Ribitsch, D.; Gübitz, G. M.; Seno, S.; Uchihashi, T.; Onoda, A. Enhanced Adsorption and Enzymatic Hydrolysis of Polyethylene Terephthalate by Cutinase with an N-Terminal Hydrophobic Tether. ACS Sustainable Chem. Eng. 2025, 13 (42), 17846–17855. DOI: 10.1021/acssuschemeng.5c05212.

(33) Shuvo, M. S. R.; Ribitsch, D.; Güebitz, G. M.; Seno, S.; Onoda, A. N-Terminal modification of aromatic tether in cutinase enhances degradation of polyethylene terephthalate powder and fabrics. Chem. Lett. 2025, 54 (7), upaf121. DOI: 10.1093/chemle/upaf121.

(34) Li, Z.; Zhao, R.; Wu, X.; Sun, Y.; Yao, M.; Li, J.; Xu, Y.; Gu, J. Identification and characterization of a novel peptide ligand of epidermal growth factor receptor for targeted delivery of therapeutics. FASEB J. 2005, 19 (14), 1978–1985. DOI: 10.1096/fj.05-4058com.

(35) Brinkman, A. M.; Chen, G.; Wang, Y.; Hedman, C. J.; Sherer, N. M.; Havighurst, T. C.; Gong, S.; Xu, W. Aminoflavone-loaded EGFR-targeted unimolecular micelle nanoparticles exhibit anti-cancer effects in triple negative breast cancer. Biomaterials 2016, 101, 20–31. DOI: 10.1016/j.biomaterials.2016.05.041.

(36) Chen, G.; Wang, Y.; Xie, R.; Gong, S. Tumor-targeted pH/redox dual-sensitive unimolecular nanoparticles for efficient siRNA delivery. J. Controlled Release 2017, 259, 105–114. DOI: 10.1016/j.jconrel.2017.01.042.

(37) Cheng, L.; Huang, F. Z.; Cheng, L. F.; Zhu, Y. Q.; Hu, Q.; Li, L.; Wei, L.; Chen, D. W. GE11-modified liposomes for non-small cell lung cancer targeting: preparation, ex vitro and in vivo evaluation. Int. J. Nanomed. 2014, 9, 921–935. DOI: 10.2147/IJN.S53310.

(38) de Paiva, I. M.; Vakili, M. R.; Soleimani, A. H.; Tabatabaei Dakhili, S. A.; Munira, S.; Paladino, M.; Martin, G.; Jirik, F. R.; Hall, D. G.; Weinfeld, M.;, et al. Biodistribution and Activity of EGFR Targeted Polymeric Micelles Delivering a New Inhibitor of DNA Repair to Orthotopic Colorectal Cancer Xenografts with Metastasis. *Mol*. Pharmaceutics 2022, 19 (6), 1825–1838. DOI: 10.1021/acs.molpharmaceut.1c00918.

